# Lysis, Lysogeny, and Virus-Microbe Ratios

**DOI:** 10.1101/051805

**Authors:** Joshua S. Weitz, Stephen J. Beckett, Jennifer R. Brum, B. B. Cael, Jonathan Dushoff

**Author notes:** Electronic address (Joshua S. Weitz).

## Abstract

We show that neither the Piggyback-the-Winner model nor coral reef virome data presented in Knowles et al. [1] support a mechanistic link between increases in lysogeny, suppression of lysis, and the decline of the virus-to-microbial cell ratio (VMR) at high microbial cell densities across environmental and human-associated systems.

---

In 1989, Bergh and colleagues used a culture-independent approach to show that virus densities in aquatic environments are thousands to millions times higher than culture-based estimates [2]. Since then, estimating the relative abundance of viruses and microbial cells has become central to efforts to characterize the scope of virus effects on ecosystem function [3–5]. A consensus had emerged from these estimates: viruses are typically 10-fold more abundant than microbial cells [6–10]. Two recent papers – by Wigington et al. [11] and Knowles et al. [1] – re-examined the collective evidence underlying this consensus in light of acknowledged variability in abundances. Using complementary datasets and analysis methods they each found that the virus to microbial cell ratio (VMR) exhibits significant variability and is *not* well described with a 10:1 or any other fixed ratio. Instead, the VMR is better described as a sublinear, power-law relationship in which the VMR decreases with increasing cell density. An equivalent interpretation is that systems with higher microbial densities tend to have more viruses in total but fewer viruses per-microbe.

Knowles et al. [1] take a further step by offering a mechanistic explanation for the emergent, nonlinear relationship. Their Piggyback-the-Winner (PtW) model states that “lytic dynamics are suppressed at high host density and density-dependent growth rate owing to the increased prevalence of lysogeny (modeled as lower specific viral production rates per infection) and super-infection exclusion rather than resistance.” They argue that this suppression of lytic dynamics due to increased lysogeny with increasing microbial density leads to the observed nonlinear relationship between viruses and microbial cell densities. In doing so, Knowles et al. [1] also claim to exclude a set of alternative explanations, including variations on Lotka-Volterra dynamics [12] and Kill-the-Winner models [7]. This exclusion is based on theoretical grounds and suggests that only PtW can explain the observed, nonlinear relationships.

We agree with Knowles et al. [1] that empirical evidence supports a systematic decline in VMR with increasing microbial cell density (see related work in [11]). However, we disagree that there is clear evidence that declining VMR is driven by a systematic increase in lysogeny. Here, we show that the PtW model does not explicitly include lysogeny. Instead, the PtW model considers lysis and lysogeny in an inconsistent manner so that lysis rates and viral release rates have distinct functional dependencies on host density. As a consequence, the PtW model exhibits enhanced lysis with increasing cell density. In addition, we find that other simple, dynamic models that do not include lysogeny predict a decline of VMR with increasing microbial cell density (see Figure 1). The rationale is that many dynamical models of virus-microbe interactions can yield nonlinear relationships between virus and microbial cell density when parameter variation is considered.

**FIG. 1:**
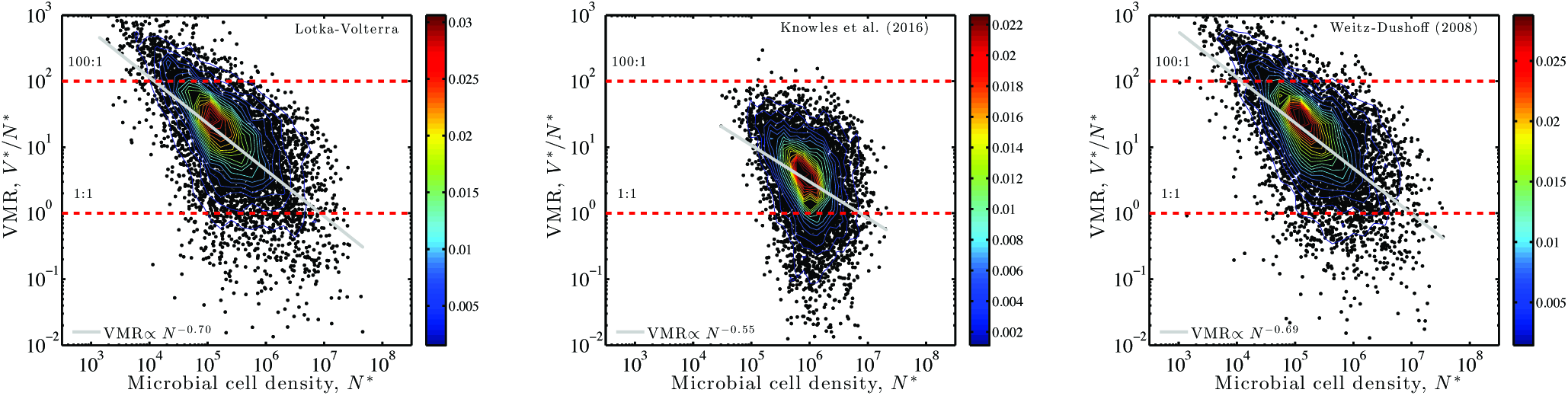
Virus density is nonlinearly related to microbial cell density in multiple dynamic models, with declining VMR as a function of increasing microbial cell density. Models include (left) Lotka-Volterra; (middle) Lotka-Volterra with density-enhanced burst sizes, aka the Piggyback-the-Winner model; (right) Lotka-Volterra with growth-limited infection and lysis. The panels show variation in model-generated VMR (black points), associated contours (colored lines, with colorbar), best fit power-law (gray line), and 1:1 and 100:1 relationships for context. See Methods for more details.

The absence of an explicit mechanistic treatment of the “prevalence of lysogeny” in the PtW model carries over to Knowles et al. [1] analysis of viromic data. The “prevalence of lysogeny” is estimated using signals derived only from viral metagenomes, rather than from cellular metagenomes. As such, measured signals conflate changes in the fraction of lysogens, the fraction of temperate phage, and the rate of lysogen induction and lysis. Increases in cell lysis by temperate phage is also consistent with their data – not all of which are statistically significant as originally claimed (see Table I) – and yet such a mechanism is in direct opposition to the notion of PtW. Indeed, measurements taken in aquatic systems support a contrasting model in which the lysogenic cell fraction decreases with increasing cell density [13].

**TABLE I:**
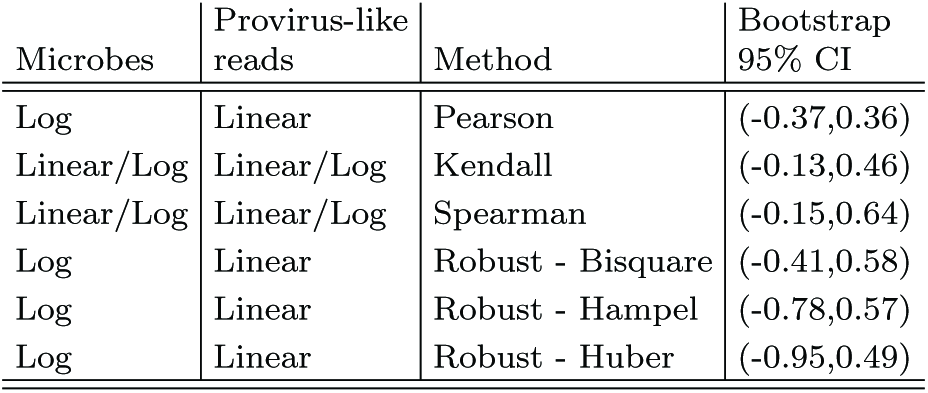
No evidence for a significant, explanatory relationship between microbial cell density and provirus-like reads in coral reefs using bootstrap methods. Each row denotes a distinct method. The 95% CIs are derived from 10,000 bootstrap attempts to evaluate the correlation using Pearson, Kendall, and Spearman metrics, and to evaluate the regression slope using three robust regression methods using alternative weighting functions. All relationships are non-significant, i.e., their 95% CIs include both positive and negative signs. Original data from Extended Data Figure 4 [1].

| Microbes | Provirus-like reads | Method | Bootstrap 95% CI |
| --- | --- | --- | --- |
| Log | Linear | Pearson | (-0.37,0.36) |
| Linear/Log | Linear/Log | Kendall | (-0.13,0.46) |
| Linear/Log | Linear/Log | Spearman | (-0.15,0.64) |
| Log | Linear | Robust - Bisquare | (-0.41,0.58) |
| Log | Linear | Robust - Hampel | (-0.78,0.57) |
| Log | Linear | Robust - Huber | (-0.95,0.49) |

In closing, given the re-analysis presented here, we urge the community not to prematurely exclude potential mechanisms in efforts to explain the observed decline of VMR with increasing microbial cell density. We are optimistic that greater consideration of interaction modes including lysis and lysogeny, spatially explicit dynamics, and variation in quantitative life history traits will shed light on the relevance of viruses to the turnover of microbial cells and nutrients on a global scale.

## Re-examining the link between dynamic models of virus-microbe interactions and VMR

We revisit the claims of what dynamic models of virus-microbe interactions can, cannot, and do say with respect to the scaling of virus density with microbial cell density. We begin with a standard model of lytic virus-microbe interactions

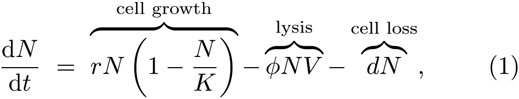

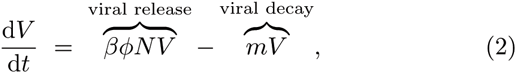

where *N* and *V* denote densities of microbial hosts and viruses, respectively. In this Lotka-Volterra model, the steady state densities are top-down controlled:

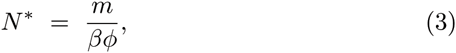

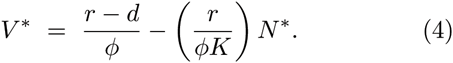

Inclusion of delays between infection and lysis need not qualitatively change the steady state values though delays can induce oscillations [14]. The virus density, *V\**, and microbial cell density, *N\**, appear to always be *negatively* related with a coefficient *r*/(*ϕK*), in contrast to the positive relationship observed in aggregated datasets. This is not the case.

Instead, viral and microbial cell densities at steady state are determined by the combined effect of multiple life history traits, which may themselves be influenced by exogenous factors. To illustrate this point, it is important to write the steady state densities as

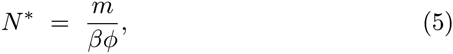

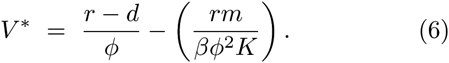

There are three parameters in the Lotka-Volterra model that could change host density at steady state: *m* - the virus decay rate, *β* - the effective burst size, and *ϕ* - the adsorption rate. For example, systems with lower adsorption rates are predicted to have higher host densities. Decreasing adsorption leads to fewer successful infection events, and an increase in host cells. A larger number of host cells supports more viruses, despite the fact that they are less efficient at adsorption, at least to a point. So long as the viral-controlled host cell density remains sufficiently smaller than the resource-limited cell density, then virus and microbial host densities are *positively* related. The key point is that *N\** is not an independently tuned variable. The steady state densities *N\** and *V\** in this model result from top-down control of hosts by viruses.

Knowles et al. [1] assume that the exogenous “carrying capacity” in the absence of viruses governs variation in both host and virus densities. Yet, neither the host-growth saturation density, *K*, or the adsorption rate, *ϕ*, are likely to be the exclusive factor varying across sample sites. What type of relationship is expected amongst virus and microbial densities if all parameters in the Lotka-Volterra model are allowed to vary? To answer this question we utilized a Latin-Hypercube Scheme (LHS) [15] to explore the effects of parameter variation on VMR (see Methods). The resulting power-law fit of virus and microbial densities is *V* ϕ *N*^0.3^. Put simply: a Lotka-Volterra model leads to nonlinear relationships between virus and microbial cell density when parameter variation is considered. As a consequence, the VMR declines with microbial cell density (see Figure 1-left) from nearly 100 to 1 while *N\** varies between 10^4^ to 10^7^ cells/ml. We also note that there is significant scatter in the model output. Aggregated empirical datasets also exhibit sublinear scaling and relatively poor explanatory power [11].

The PtW model is intended to reconcile the inability of alternative, simple models to explain the decrease of VMR with increasing microbial cell density. To do so, Knowles et al. [1] claim that lysis in the PtW model is suppressed at high densities due to the increased prevalence of lysogeny. Virus and microbial cell dynamics in the PtW model are governed by the following equations:

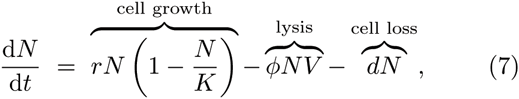

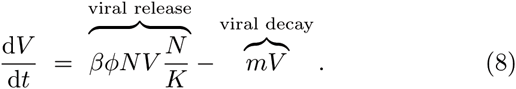

As is evident, there is no explicit class of lysogens in this model nor any induction of temperate phage despite the claim that the equations represent a model of lysogeny (see [16, 17] for examples of models that do include explicit treatment of lysogeny).

In effect, the PtW model is a variant of Lotka-Volterra models in which the burst size of lytic viruses is modulated by the relative population size, *N*/*K*. When host densities, *N*, approach *K* then burst sizes reach their maximum, whereas when host densities, *N*, are much less than *K* then burst sizes approach 0. Hence in PtW, a valid interpretation is that burst size is enhanced at high cell densities and suppressed at low cell densities. Alternatively, the claim that lysis is suppressed at high host densities could be interpreted to mean that the relative burst size per lysed cell decreases with increasing *K*. However, doing so conflates the death of cells via lysis with viral release.

In contrast, one may ask: what is the relative lysis rate when comparing low and high densities due to variation driven exclusively by changes in *K*? In the PtW model the lysis of microbial cells has the same functional form as it does in the Lotka-Volterra model. Both host and viral steady state densities increase with *K* in the PtW model. As a consequence the total lysis rate (*ϕN* V\**) and per-capita lysis rate (*ϕV\**) increase with increasing *K* at steady state. These relationships are inconsistent with the claim that lysis is suppressed in the PtW model at high host densities. Nonetheless, we apply a LHS approach to evaluate the relationship between VMR and microbial cell density given variation in life history traits (see Methods). In the PtW model, as in the Lotka-Volterra model, viral and microbial cell densities are non-linearly related, i.e., *V* α *N*^0.45^, such that VMR declines with increasing cell density (Figure 1-middle).

How general is this qualitative finding? One of the simple models that Knowles et al. [1] critique is another variant of Lotka-Volterra dynamics proposed by Weitz and Dushoff [12]. The dynamics are:

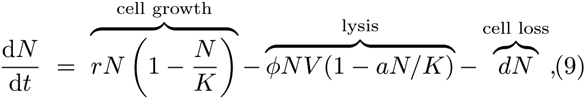

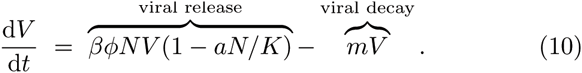

The original purpose of this model was to assess how implicit resource limitation would affect virus-host dynamics. Viruses require cellular machinery to reproduce, therefore the model assumes that when cell growth slows then viral infection and lysis should also slow. As a consequence, it is possible for slow-growing populations of bacteria to avoid top-down control by viruses. This model has been extended to resource-explicit dynamics with similar conclusions [17]. This model is equivalent to a standard Lotka Volterra model when a = 0 [12] as is self-evident from inspection of the equations reproduced here. We assess the relationship between *V\** and *N\** given a LHS analysis. We find that the best-fit power-law model is *V* α *N*^0.31^ and that VMR declines with increasing microbial cell density (Figure 1-right).

We remain cautious with respect to declaring victory prematurely, despite the “agreement” between each of these dynamic models and the empirical finding of a declining VMR with microbial cell density [1, 11]. Models of virus-microbe dynamics include various degrees of complexity or realism - depending on one’s perspective. Simple models can often yield complex, emergent behavior, whereas complex models can often yield simple behavior. The choice of modeling details reflects the fact that these models are meant to be used for different problems, i.e., in the spirit of P.W. Anderson’s adage “more is different” [18].

The three models assessed here describe a single population, or functional group, of viruses interacting with a single population, or functional group, of microbial hosts. None of the three models explicitly accounts for stochastic dynamics, multiple strains, impact of higher trophic levels, or even lysogeny. The effects of many such extensions are treated at length in [14]. These extensions may be compatible with nonlinear scaling of *V\** with *N\** (see [19] for one such example, in which Lotka-Volterra models are embedded as part of a complex, multi-strain model). Would such agreement alone constitute proof of the correctness of the model(s)? We contend that the appropriate answer to this question is: no. Rather, such an agreement would point to the need to test both model assumptions and output. Similar critiques have been raised with respect to interpreting the utility of models in cellular systems biology [20].

In the long term, identifying models that “fit” large-scale patterns can help expand the potential set of mechanisms that are compatible with observations. An expanded set of mechanisms can help motivate the research community to revisit old assumptions or test new ones. For over 25 years, the environmental virology community has measured and analyzed total virus and microbial cell abundances [1–5, 11, 21]. In addition to understanding the basis for variation in such abundances, it is essential to probe the functional scale of interactions amongst a network of viruses and microbial hosts [22, 23]. These interactions are likely to include lysogeny and other modes of interactions. In doing so, it is imperative to consider how mechanisms of virus-host interactions assumed by dynamic models at cellular scales translate into population-and global-scale consequences.

## Re-examining the relationship between the “prevalence of lysogeny” and microbial cell densities

The core mechanism proposed in the PtW model is that of suppressed lysis due to an increasing importance of lysogeny at high microbial cell densities. What is the statistical evidence that the prevalence of lysogeny increases across sites where virus and microbial cell densities were collected? Knowles et al. [1] do not have any direct measurements for the bulk of their sites because evidence for lysogeny is collected exclusively within coral reefs. Instead, the lysogenic prevalence in coral reef viromes is measured in terms of three proxies: the frequency of integrase, excisionase, and provirus-like reads. The use of viromes as the exclusive means for estimating the prevalence of lysogeny in coral reef ecosystems raises a potential dilemma. The life history of temperate phage includes both an intracellular and extracellular stage. When lysis is suppressed, temperate phage should preferentially remain integrated inside their host genomes as prophage. If fewer temperate phage induce and lyse their host cells then there should be a decrease in abundance of provirus-like reads and other proxies for lysogenic prevalence as measured in viromes. This is not the interpretation of Knowles et al. [1]. Rather they interpret an increase in provirus-like reads and related proxies to indicate an increase in the prevalence of lysogeny.

For the moment, we adopt the perspective of Knowles et al. [1] and re-examine the relationship between provirus-like reads in coral reef viromes and microbial cell densities (Extended Data Figure 4, [1]). Knowles et al. [1] utilize “robust regression” techniques to find a significant, positive relationship between provirus-like reads and microbial cell densities. In contrast, we find no statistically significant relationship between provirus-like reads and microbial cell densities when evaluated using correlations-based and robust regression approaches (Table I). The fraction of integrase and excisionase in coral reef viromes represent the other purported signals of lysogeny prevalence. Knowles et al. [1] find 90%, rather than 95%, CI support for a positive relationship between integrase and microbial cell density. They also find 95% CI support for a positive relationship between excisionase and microbial cell density.

We contend there is an alternative interpretation to the finding that one proxy, i.e., excisionase, rather than three proxies for lysogeny prevalence have a significant positive relationship with microbial cell density with at least 95% CI support. A shift of temperate phage to the lytic mode at increasing microbial cell densities could lead to elevated levels of temperate phage hallmark genes detected in viromes from extracellular viral communities. This interpretation is diametrically opposed to the conception of PtW that lysis is suppressed at high microbial cell densities. If Knowles et al. [1] had measured a provirus-like signal within cellular metagenomes, that could have provided additional clarifying information. For example, a testable hypothesis of a mechanistic adaptation of PtW is that the fraction of lysogenic cells, i.e., cells with integrated prophage, increases with increasing microbial cell density. The absence of complementary evidence to disentangle the fraction of temperate phage within the viral community, the fraction of lysogenic cells, and the rate at which prophage induce cells prevents definitive conclusions regarding the role and relevance of lysogeny in coral reef ecosystems.

We conclude that there is little empirical evidence to support the claim of Knowles et al. [1] that lysis is suppressed at high microbial cell densities within coral reef ecosystems due to an increase in lysogeny prevalence. In contrast, measurements in aquatic systems support a contrasting model in which the fraction of lysogenic cells decreases with increasing microbial cell density [13]. This direct evidence is partially confirmed in Knowles et al. [1] (Extended Data Figure 2) when individual studies are examined prior to aggregation. It remains an open question as to the generality of this finding. Further work is needed to characterize the relative abundance and role of lysogeny in shaping dynamics within environmental and human-associated systems.

## Methods

### Dynamic model analysis

We use baseline parameter values *r* = 1/24 hrs^−1^, *d* = 1/48 hrs^−1^, *K* = 10^7^ cells/ml, *ϕ* = 10^−8^ ml/cells/hr, *m* = 1/6 hrs^−1^, and *β* = 20 for the analysis of the Lotka-Volterra, *β* = 20.2 and *a* = 0.1 for the Weitz-Dushoff [12] models and *β* = 200 for the analysis of Knowles et al. [1]. In this way the effective burst size is 20 for all models when *N* = *K*/10.

For the LHS analysis implemented in MATLAB we permit each parameter to vary ten-fold above and below. We generated 10^4^ random parameter combinations and analyzed only those that led to coexistence.

### Bootstrap analysis of indicators of lysogeny

We implemented a bootstrap analysis using R to identify 95% confidence intervals (CIs) for the relationship between microbial cell density and the percentage of provirus-like reads in coral viromes. Following Knowles et al. [1], we define the *x*-variable to be the log_10_ transformation of microbial cell density, measured in units of counts per ml, and the *y*-variable to be the percentage of provirus-like reads. We used these variable definitions for assessing the significance of correlations via a parametric approach – Pearson moment-product – and two non-parametric approaches – Spearman rank and Kendall’s tau. The non-parametric correlations are insensitive to transformations of the microbial cell density. Similarly, we used the same variable definitions in a robust regression analysis to estimate the slope of the relationship using alternative weighting functions that weight “outliers” differently. A total of 10^4^ bootstrap replicate attempts were made in each case.

### Code

All scripts are available on github at https://github.com/WeitzGroup/VMR-Lysis-Lysogeny.

